# Joint modelling of environmental and species responses to restoration

**DOI:** 10.1101/2025.03.24.644885

**Authors:** Aapo Jantunen, Otso Ovaskainen, Atte Komonen, Merja Elo

## Abstract

Restoration outcomes have unexplained variation. As environmental variables can both affect the restoration outcome and be affected by restoration, studying the interplay between environmental and species responses could reveal reasons behind this variation. Here, we present a methodological framework of jointly modelling environmental and species responses to restoration and demonstrate it by studying the effect of water table (WT) on vegetation in restored forestry-drained boreal peatlands. Restoration was consistently successful in raising WT in the higher but not in the lower productivity peatland types. Jointly modeling WT and vegetation produced the highest accuracy for predicting vegetation response, highlighting the need to monitor relevant environmental factors in restoration research. The study demonstrates how the framework can be used to explain differences in restoration outcomes, even with sparse environmental data. The framework allows predicting restoration outcomes in different scenarios of environmental variables, facilitating the development of restoration methods with predictable outcomes.

**Implications for practice:**

- Monitoring environmental factors that could affect targeted taxa is important in restoration research
- Jointly modelling environmental and species responses to restoration can reveal factors behind variation in restoration outcomes

## Introduction

Ecosystem restoration is increasing globally due to agreements such as Kunming-Montreal Global Biodiversity Framework (CBD 2022) and European Union’s Nature Restoration Law (European Union 2024). Restoration is effective in improving ecosystem services (Benayas et al. 2009) and biodiversity (Atkinson et al. 2022) of degraded ecosystems. However, restoration outcomes vary between and within ecosystems (Atkinson et al. 2022).

One common aim of restoration is to initiate the recovery of species communities in degraded sites towards those in pristine sites (Suding 2011). In many cases, restoration actions do not target the species directly. Instead, they target the abiotic or environmental factors that shape the species communities. This approach is called the Field of Dreams hypothesis (Palmer et al. 1997). Its idea leans on two of the four processes of community ecology: dispersal and selection (Vellend 2008). The targeted species must be able to disperse to the restored sites, and their fitness must be higher than that of current species.

To increase predictability of restoration, it is important to understand what factors affect the restoration outcome and its variation (Brudvig et al. 2017). We need methods to understand why restoration was successful in one site, but not in the other. For this purpose, we present an analytical framework of jointly modelling the environmental and species responses to restoration. With this method, we answer how much of the variation in species responses are explained by the environmental factor that was the target of restoration actions. We show how examining the co-variation between environmental and species responses can reveal factors behind variation in restoration outcomes.

We use data from a replicated long-term before-after control-impact study of forestry-drained boreal peatlands (Elo et al. 2024a; b). In general, peatlands are restored for biodiversity (Minayeva et al. 2017) and different ecosystem services such as carbon sequestration (Wilson et al. 2016). As WT is an important factor that shapes the species communities in peatlands, peatland restoration usually begins with raising the WT (Andersen et al. 2016). In forestry-drained peatlands, this is achieved by blocking and/or damming ditches and cutting down trees to reduce water flow and decrease evapotranspiration (Kuuluvainen et al. 2002). The rise in WT starts succession from communities of forest and generalist species towards communities found in pristine peatlands (Jauhiainen et al. 2002, Haapalehto et al. 2017).

Despite similar restoration methods, Elo et al. (2024a) found considerable variation in restoration outcomes. Some of the variation was caused by peatland type, but the effect of WT on the variation was not studied (Elo et al. 2024a). Here, we explain the variation Elo et al. (2024a) observed with unpublished sparse plot level WT data.

## Methods

### Data gathering

The study consists of 101 sites in southern boreal and northern boreal biogeographical zone in Finland (please see the full description of the data in Elo et al. 2024a). The sites belong to four peatland types: poor pine mire forests, rich pine mire forests, poor open mires, and rich open mires. Pine mire forests have Scots pine (*Pinus sylvestris*), while open mires have little to no trees. They are dominated by peat-forming *Sphagnum* mosses, with different dwarf shrubs, particularly in pine mire forests, and sedges, such as *Carex*, particularly in open mires. In both pine mire forests and open mires, the poor sites are ombrotrophic and rich sites vary from oligotrophic to mesotrophic.

Each peatland type has sites in three different treatments: drained (4–6 sites per type), restored (9–11 sites), and pristine (9–11 sites). Drained and restored sites were drained for forestry in the 1960s and 1970s. Restored sites were restored between 2007 and 2014 to raise WT by filling the ditches with peat and building dams to block water flow and cutting down trees to reduce transpiration and shadowing. Pristine sites have not been drained.

All sites were monitored for vegetation and WT. Vegetation monitoring was conducted on ten permanent 1 m x 1 m plots (Figure S1), from which cover (%) of each vascular plant and moss was visually estimated. The location of the monitoring plot complex was randomly chosen in a place that represented typical vegetation in the site. In drained and restored sites, the distances from plots to ditches were at least ten meters. Plots were four meters away from each other in two rows. WT was monitored from wells (perforated hollow plastic pipes) in one or two corners of each vegetation plot. The first sampling was before restoration (year 0) and was repeated at years 2, 5 and 10 after restoration. Same sample interval was also used in drained and pristine sites. Sampling was done between years 2006 and 2022.

### Statistical analysis

For the analysis, we selected three species groups and two individual species (‘taxa’ from now on), which have shown to have strong responses to restoration (Elo et al. 2024a). The species groups were *Sphagnum* mosses (32 species) and *Carex* (23) species, which increase after restoration, and common forest mosses (*Dicranum polysetum, Hylocomium splendens, Pleurozium schreberi*), which decrease after restoration. The main habitat for the forest mosses is either heath forests or mesic and herb-rich heath forests (Finnish Biodiversity Information Facility 2024). In addition, we selected birch (*Betula pubescens*) and tussock cottongrass (*Eriophorum vaginatum*), which both increase after restoration. The list of the 60 included species is in Table S1.

WT data were relatively sparse: we had it for 68 % of the datapoints (time x plot), and 28 % of the plots had all four measurements. We divided WT to two variables: whether it is above -40 cm (binary, 1/ 0; ‘HIGH’) and if it is, the measure of the WT (continuous, -40–+20 cm; conditional water table ‘CWT’). Negative CWT means that the WT is below peat surface. With these two variables, we retained information of dry wells where the WT was very low, but the exact value was unknown. We chose - 40 cm as the cut off value based on the estimated WT tolerance of bryophytes estimated by Menberu et al. (2016). We discarded values higher than +20 cm as they were likely due to rainwater accumulation in the well. Such high values were not observed when WT was monitored with automated loggers from a subset of the sites (Menberu et al. 2016). When the WT was measured from two wells in one vegetation plot, we used their mean.

We used Hierarchical Modelling of Species Communities to model responses of WT and vegetation to restoration (Ovaskainen et al. 2017). Response variables were HIGH, CWT, occurrence of each taxon and cover of each taxon. We used probit model for HIGH and occurrence, and normal model for CWT and cover. As CWT is defined only if HIGH is true, and cover is defined only if the species is present, these variables had missing entries. Cover was log-transformed. CWT and cover were normalized to zero mean and unit variance. Fixed part of the model consisted of treatment (three-level factor: drained, restored, pristine), time (continuous variable: 0, 2, 5, 10 years), time squared, interaction of treatment and time, and interaction of treatment and time squared. We included year of the sampling, plot number, and location of the peatland as random effects modelled through latent variables (Ovaskainen et al. 2017). We created separate models for each peatland type. We could not model the cover of *Betula pubescens* in poor open mires, as its occurrence was so low.

Hierarchical Modelling of Species Communities uses Bayesian inference through Markov chain Monte Carlo sampling (MCMC) (Tikhonov et al. 2020). We sampled the posterior distribution with four chains of 250 samples with thinning of 1 000 each. We used 500 000 steps for the transient and adaptation phase. We used default priors (Tikhonov et al. 2020). To examine if parameter convergence was satisfactory, we used effective sample size (Figure S2) and potential scale reduction factor (Figure S3) of the posterior sample. We used explanatory power to assess model fit, using Tjur’s R^2^ for binary and R^2^ for continuous variables. We did the modelling in R (4.3.2) and used the package Hmsc (3.1-3, Tikhonov et al. 2020).

We calculated taxon abundance as a product of probability of presence and cover when the taxon is present. We calculated restoration effect as the difference of change between restored and drained sites for WT variables and taxon abundances but as vegetational changes have already been studied (Elo et al. 2024a), we will not discuss them further. We report medians of the 1 000 MCMC-samples and consider the median having high and moderate statistical support if 95 % and 80 %, respectively, of the samples showed same sign as median.

Restoration effect = (Var_r, 10_ – Var _r, 0_) – (Var_d, 10_ – Var_d, 0_)

where Var means variable (HIGH, CWT or taxon abundance), r means restored, and d means drained. 10 and 0 refer to the time of sampling.

We used species association matrices to study the effect of WT on the taxa. These matrices show the residual correlations (correlations that are not explained by the fixed part of the model) between all response variables. We focused on site-level associations.

We used four-fold cross-validation and conditional cross-validation to see whether WT data improved predictive power of the taxa models. As cross-validation uses substantial computational resources, we computed them with models using thinning of 100 instead of 1 000. Their convergence was slightly worse (Figure S2, Figure S3).

To visualize the effect of WT, we calculated predicted abundances of taxa in six different WT scenarios (unknown, <-40 cm, -35 cm, -25 cm, -15 cm, -5 cm). We calculated predictions for those taxon-peatland type pairs, where one of the WT variables had high support association with taxon abundance and the same WT variable improved predictive power by at least 0.05. We calculated these predictions for restored sites 10 years after restoration. We selected one restored site for each peatland type, vegetation plot 10, year 2018 and assumed there were no spatial random effects. We made these choices to make general predictions where WT was the only factor with variation. We also tested other values to confirm that the effect of WT stayed similar in different scenarios.

From the total of six peatland ecosystem types included in the original vegetation data (Elo et al. (2024), we selected four. We excluded spruce mire forests because we could not use conditional cross-validation with them due enduring errors in MCMC sampling. These errors were most probably caused by their high heterogeneity, which is underlined when only a subsample of the sites is used as training data.

## Results

### Restoration effect in WT and taxa abundances

Restoration raised the mean WT in rich pine mire forests and rich open mires (Figure 1, Figure 2). By contrast, restoration effect on WT was unnoticeable in poor pine mire forests and poor open mires. However, the difference between pristine and restored sites diminished in poor pine mire forests and poor open mires due to decrease in WT in pristine sites (Table S2). There was also moderate support for restoration effect in WT in poor open mires in the first five years, after which the WT declined (Figure 1, Table S2). Sites selected for restoration were not identical to drained controls at year 0. There was high support for higher WT in both poor and rich pine mire forests compared to drained sites in both WT variables (Figure 1, Table S2).

**Figure 1.**
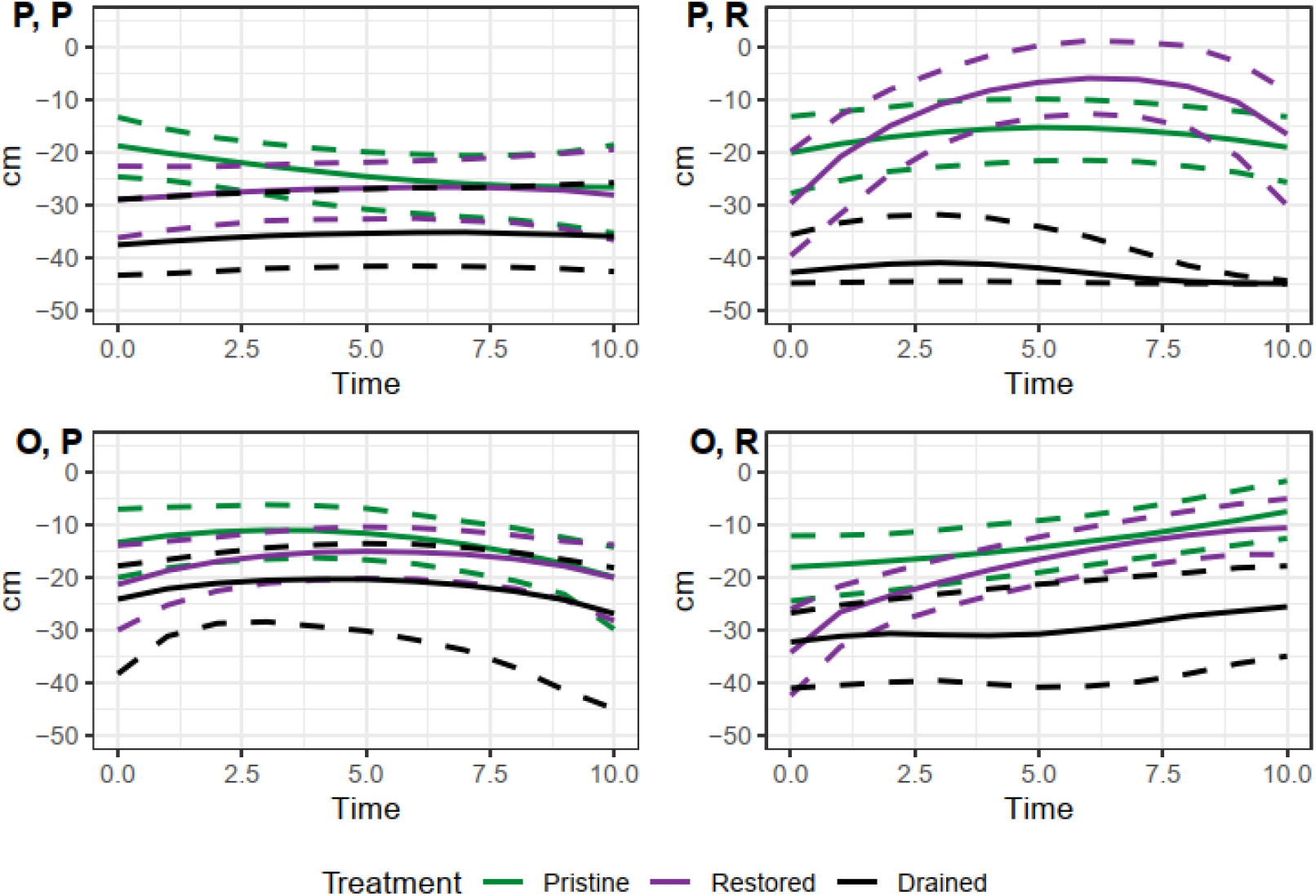
Change of water table level in pristine, restored and drained treatments in different peatlxand types. Lines show the median of 1 000 MCMC samples. 90 % of the samples were between dashed lines. Probability of water table level being over - 40 cm (HIGH) and exact water table level, if it was (CWTL), are combined to a single value to visualize changes in water table level. If water table level was below -40 cm, it was set to -45 cm. If it was above -40 cm, then the exact value was used. Abbreviations: P: pine mire forests, O: open mires. P: poor, R: rich.

**Figure 2.**
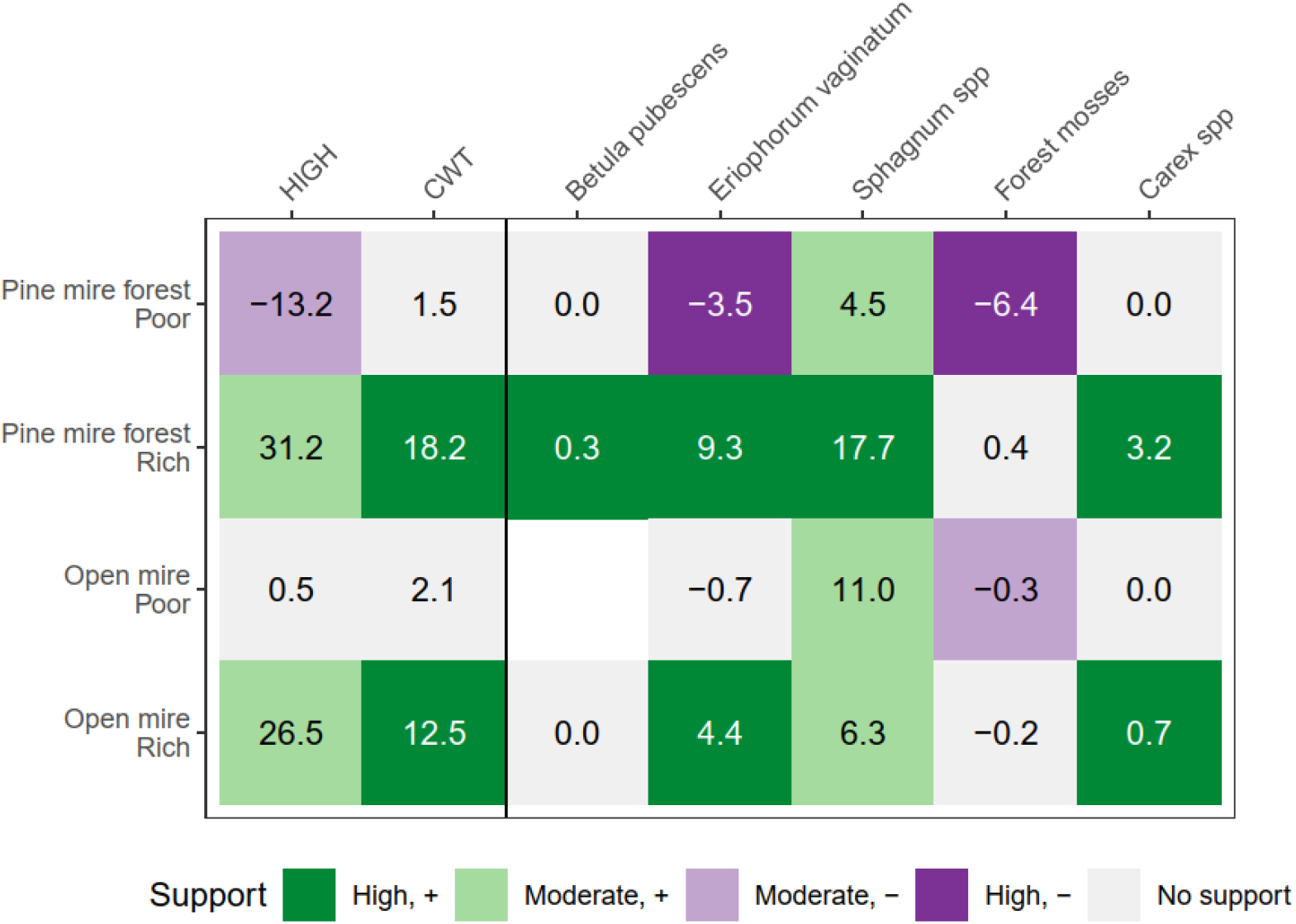
Median restoration effects in water table level variables and taxon abundances in different peatland types. HIGH refers to probability of water table level reaching higher than -40 cm. CWT refers to exact water table level if it was higher than -40 cm. Abundance of *Betula pubescens* could not be modelled in poor open mires due to low occurrence. Support levels: High: 95 % of the MCMC samples have same sign as median. Moderate: 80 % of the samples have same sign as median

Vegetation responses to restoration varied between peatland types (Figure 2). Restoration effect was the strongest in rich pine mire forests where four taxa was affected with high support. Poor open mires was the only peatland type with no high support effects.

### WT as a predictor for abundance

WT improved predictive power of the taxon models in 16 out of 40 of the cases by more than 0.01 (Figure 3). The improvements were mostly small, but for example, the predictive power of occurrence model of *Eriophorum vaginatum* in poor open mires increased six-fold from 0.02 to 0.13. Forest moss occurrence in rich open mires had both largest improvement in predictive power and highest predictive power (0.22, 0.42 respectively).

**Figure 3.**
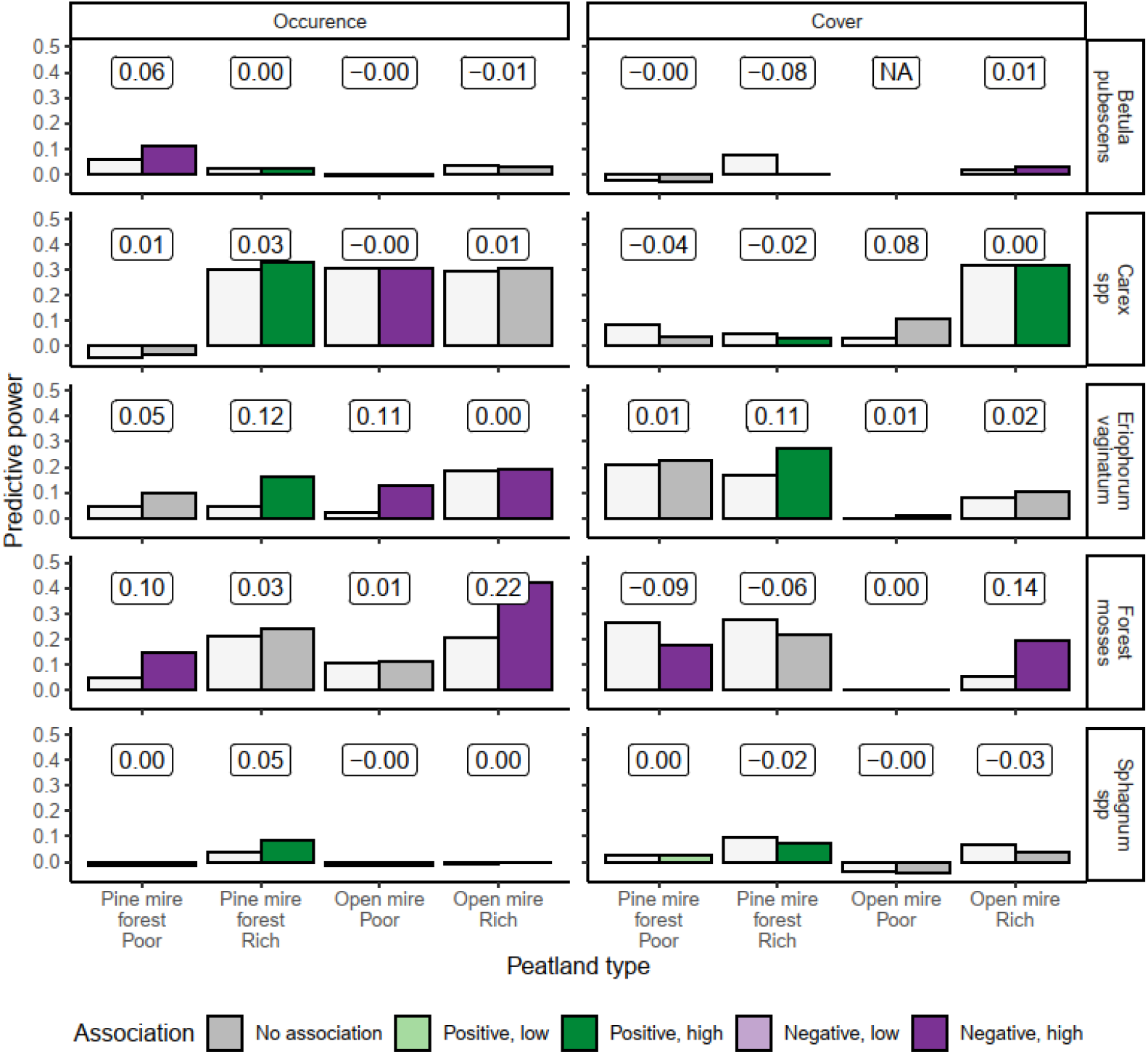
Predictive power of vegetation models when water table levels was not known (first bar) and when it was (second bar). The numbers above bars show improvement of predictive power when water table level was known. Negative values indicate decrease in predictive power. Colours of the second bar show associations of taxon and water table level calculated from models fitted with all observations (required support level: 80 %). Association is shown if either one or both of water table level variables were associated with taxon abundance. High refers to residual correlation >0.33 and low <0.33.

WT improved predictions more frequently for taxon occurrence models (11 of 20 models) than for cover models (5 of 19 models). Similarly, predictive power decreased in none of the occurrence models but in 7 cover models.

Occurrences of *Sphagnum* mosses and *Betula pubescens* had a very low predictive power in all peatland types (Figure 3). This is because Tjur’s R^2^ is the coefficient of discrimination, and both taxa have really unbalanced occurrence distributions. *Sphagnum* mosses are found in almost all vegetation plots in pine mire forests and open mires (>97 %), and *Betula pubescens* is very rare (<5 %). This leads to the models having similar possibility of occurrence for plots which have *Sphagnum* mosses or *Betula pubescens*, and plots which do not have them, and no discrimination between the two is achieved.

In general, HIGH affected the taxa’s abundance predictions more than CWT (Figure 4, Figure S4). But in some cases, as with the forest mosses in rich open mires, also the finer changes in CWT affect taxon abundance.

**Figure 4.**
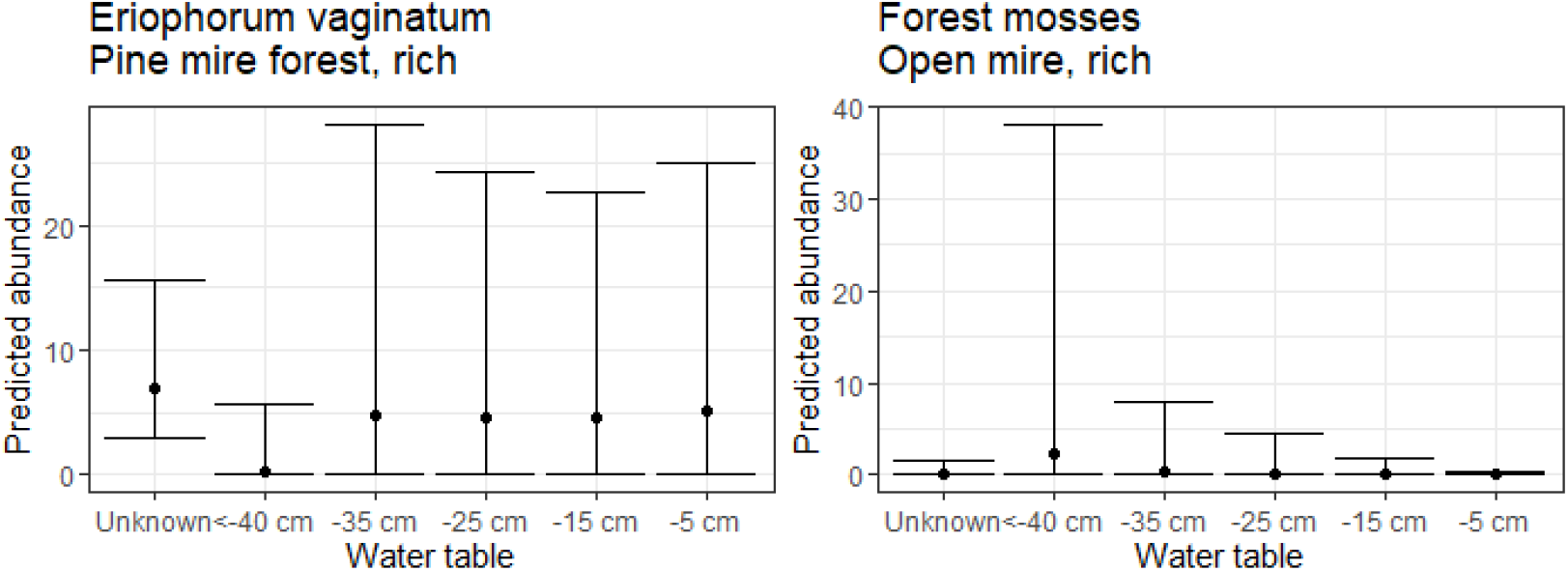
Predicted abundances of *Eriophorum vaginatum* in rich pine mires and forest mosses in rich open mires in different water table level scenarios. Predictions were calculated for restored sites 10 years after restorations. Points show median and whiskers show range between lowest and highest 10 % of the predictions of 1 000 MCMC samples.

## Discussion

We used joint species distribution models to simultaneously model taxa and environmental responses to restoration with data from restored boreal forestry-drained peatlands. We used the models to study how restoration affects the targeted environmental factor, and if this environmental factor explains differences in taxa’s responses. The results show that restoration had desired effects on WT and vegetation, but not in all peatland types. Even with the sparse environmental data, our method showed that WT explained part of the variation in restoration outcomes within peatland types. Knowledge of WT improved predictive power of all species in at least some of the peatland types.

Our goal was to understand variation in restoration outcomes between and within peatland types. While studying the same vegetation data, Elo et al. (2024) observed that restoration had smaller effect on vegetation in poor pine mire forests and poor open mires than other peatland types. They hypothesized that one cause for this could be failure in raising WT. Reasons for this could be that poor pine mire forests and poor open mires are usually located in ombrotrophic raised bogs, which makes them hydrologically isolated from the surrounding areas, and peat subsidence close to the ditches (Kuuluvainen et al. 2002). This hypothesis is in line with our observations of unnoticeable restoration effect in WT in ombrotrophic peatland types and increase in predictive power of vegetation models when WT was known.

However, the poor pine mire forests and poor open mires did not stand out with more or stronger associations between vegetation and WT as they should have, if sites with higher WT differed distinctly from sites with lower WT. This supports the other hypothesis by Elo et al. (2024a) that pre-restoration community structure might affect restoration outcome.

However, our WT measurements were sparse, with only four measurements per site in the span of ten years. WT is known to fluctuate between and within years in peatlands due to weather conditions (for example, Peichl et al. 2014). While our and previous results show that WT has clearly an effect on restoration outcomes (Hedberg et al. 2012; Maanavilja et al. 2014), with the sparse WT data at hand, we cannot quantify its full effect.

Our study corroborates previous findings in that gathering relevant environmental data in restoration studies is important part of understanding the variation in restoration outcomes (Bakker et al. 2003; Stuble et al. 2017). Same environmental factors might be affected by restoration and, in turn, affect restoration outcomes. This has been confirmed in stream restoration, where restoration affects riparian vegetation, which affects the morphological development of the stream (Vargas-Luna et al. 2018). Similar phenomena could be true with forest structure in peatland restoration. Forest structure affects species composition in restored peatlands (Maanavilja et al. 2014; Lefebvre-Ruel et al. 2018) through shadowing and increasing evapotranspiration (Hedberg et al. 2012). Restoration also affects forest structure by planned felling (Kuuluvainen et al. 2002) and increased tree mortality due to raised WT (Aapala et al. 2013).

A strength of our joint species distribution modelling method is that it can be extended to more complex cases with many environmental factors. Thus, it could be used to understand previously mentioned complex feedback systems. With this knowledge, we could develop restoration methods with predictable outcomes in a variety of ecosystem types.

## Acknowledgements

We are thankful to all the researchers, specialists, and field workers who have participated in developing and maintaining the National Peatland Restoration Monitoring Network, which has been funded by the Boreal Peatland LIFE (LIFE08 NAT/FIN/000596) and Hydrology LIFE projects (LIFE16 NAT/FIN/000583). In addition, AJ was funded by The Finnish Foundation for Nature Conservation and Kone Foundation; ME by the Finnish Ministry of Environment; OO by the Research Council of Finland (grant no. 336212 and 345110), and the European Union: the European Research Council (ERC) under the European Union’s Horizon 2020 research and innovation programme (grant agreement no. 856506; ERC-synergy project LIFEPLAN). We also thank CSC – IT Center for Science, Finland, for computational resources.

