## Supplementary material for "Joint modelling of environmental and species responses to restoration": Jantunen et al, supplementary information.docx

Figure S1. Monitoring design with ten vegetation plots, each of which had one or two water table level pipes. The black line is a line between ditches and parallel to them in restored and drained sites. Direction of the line was randomized in pristine sites.

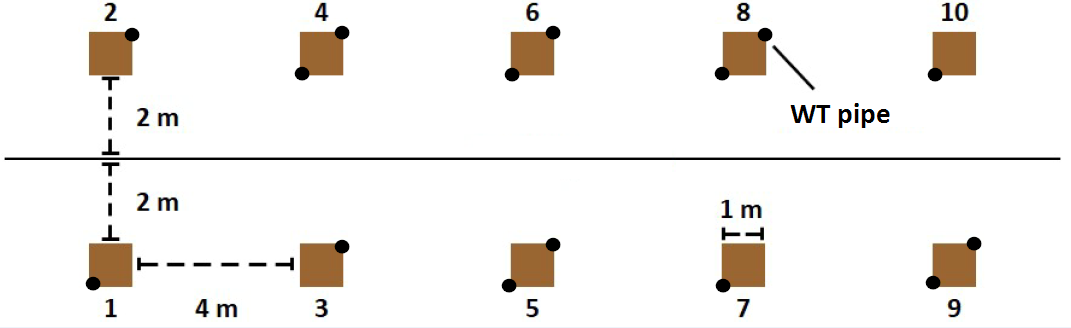

Table S1. List of studied species. Taxon refers to the species group used in the analysis.

| Taxon | Species |
| --- | --- |
| *Betula pubescens* | *Betula pubescens* |
| Carex | *Carex acuta* |
| Carex | *Carex aquatilis* |
| Carex | *Carex brunnescens* |
| Carex | *Carex buxbaumii* |
| Carex | *Carex canescens* |
| Carex | *Carex cespitosa* |
| Carex | *Carex chordorrhiza* |
| Carex | *Carex dioica* |
| Carex | *Carex disperma* |
| Carex | *Carex echinata* |
| Carex | *Carex flava* |
| Carex | *Carex globularis* |
| Carex | *Carex lasiocarpa* |
| Carex | *Carex limosa* |
| Carex | *Carex loliacea* |
| Carex | *Carex nigra* |
| Carex | *Carex pauciflora* |
| Carex | *Carex paupercula* |
| Carex | *Carex rhynchophysa* |
| Carex | *Carex rostrata* |
| Carex | *Carex rotundata* |
| Carex | *Carex vaginata* |
| Carex | *Carex vesicaria* |
| Forest mosses | *Dicranum polysetum* |
| *Eriophorum vaginatum* | *Eriophorum vaginatum* |
| Forest mosses | *Hylocomium splendens* |
| Forest mosses | *Pleurozium schreberi* |
| Sphagnum | *Sphagnum angustifolium* |
| Sphagnum | *Sphagnum annulatum* |
| Sphagnum | *Sphagnum aongstroemii* |
| Sphagnum | *Sphagnum balticum* |
| Sphagnum | *Sphagnum capillifolium* |
| Sphagnum | *Sphagnum centrale* |
| Sphagnum | *Sphagnum compactum* |
| Sphagnum | *Sphagnum cuspidatum* |
| Sphagnum | *Sphagnum fallax* |
| Sphagnum | *Sphagnum fimbriatum* |
| Sphagnum | *Sphagnum flexuosum* |
| Sphagnum | *Sphagnum fuscum* |
| Sphagnum | *Sphagnum girgensohnii* |
| Sphagnum | *Sphagnum jensenii* |
| Sphagnum | *Sphagnum lindbergii* |
| Sphagnum | *Sphagnum majus* |
| Sphagnum | *Sphagnum medium* |
| Sphagnum | *Sphagnum obtusum* |
| Sphagnum | *Sphagnum palustre* |
| Sphagnum | *Sphagnum papillosum* |
| Sphagnum | *Sphagnum platyphyllum* |
| Sphagnum | *Sphagnum pulchrum* |
| Sphagnum | *Sphagnum quinquefarium* |
| Sphagnum | *Sphagnum riparium* |
| Sphagnum | *Sphagnum rubellum* |
| Sphagnum | *Sphagnum russowii* |
| Sphagnum | *Sphagnum spp* (unspecified) |
| Sphagnum | *Sphagnum squarrosum* |
| Sphagnum | *Sphagnum subfulvum* |
| Sphagnum | *Sphagnum subnitens* |
| Sphagnum | *Sphagnum subsecundum* |
| Sphagnum | *Sphagnum tenellum* |
| Sphagnum | *Sphagnum teres* |
| Sphagnum | *Sphagnum warnstorfii* |
| Sphagnum | *Sphagnum wulfianum* |

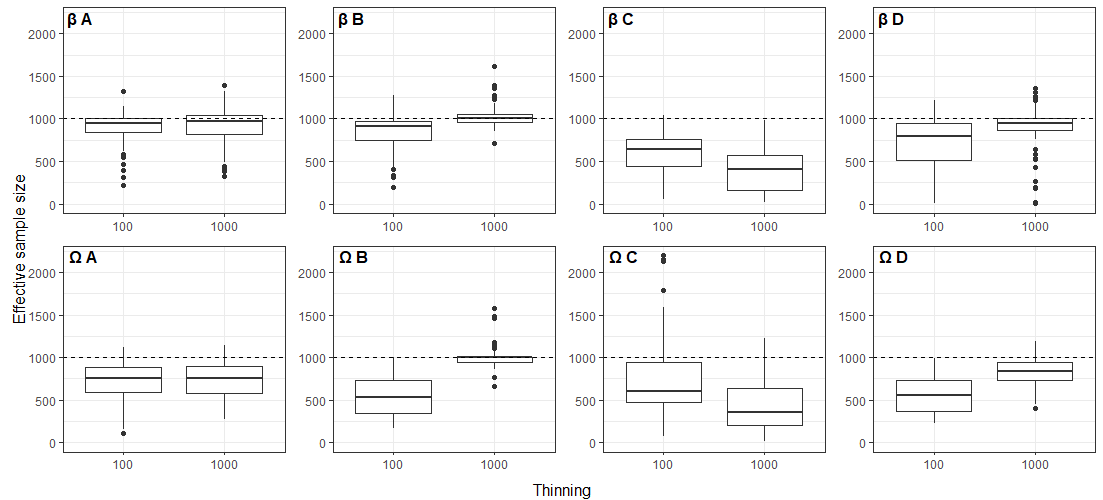

Figure S2. Effective sample sizes of Beta (β) and Omega (Ω) parameters of the models with different values of thinning parameter. Omega parameter values of species-to-species interactions were excluded. A = poor pine mire forests, B = rich pine mire forests, C = poor open mires, D = rich open mires. Four high value outliers from thinning 100 were omitted from plot ΩC for visualization.

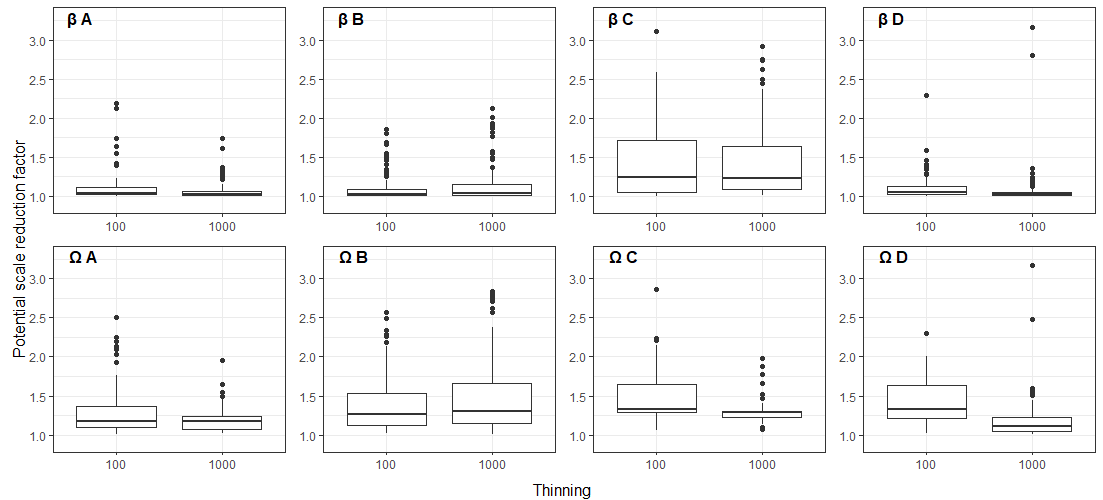

Figure S3. Potential scale reduction factors of Beta (β) and Omega (Ω) parameters of the models with different values of thinning parameter. Omega parameter values of species-to-species interactions were excluded. A = poor pine mire forests, B = rich pine mire forests, C = poor open mires, D = rich open mires. Some high value outliers were omitted for visualization: plot βC one from thinning 100 and two from thinning 1000, plot βD four from both thinning values, plot ΩC one from both thinning values.

Table S2. Changes and initial differences in water table level variables in different peatland types. Values are medians of 1 000 MCMC samples. Values inside parenthesis are proportion of samples that had positive sign for median.

| Peatland type | Variable | Restoration effect, 10 yr | Initial difference, restored vs. drained | Change, restored | Change, pristine | Change in restored vs. pristine, 10 yr | Restoration effect, 5 yr |
| --- | --- | --- | --- | --- | --- | --- | --- |
| Pine, poor | HIGH | -13.2 (0.19) | 25.5 (0.96) | -7.9 (0.25) | -13.6 (0.01) | 6.4 (0.72) | -1.0 (0.47) |
| Pine, poor | CWTL | 1.5 (0.67) | 5.7 (0.97) | 3.2 (0.81) | -4.8(0.11) | 8.2 (1.00) | -0.6 (0.43) |
| Pine, rich | HIGH | 31.2 (0.95) | 47.8 (1.00) | 13.2 (0.81) | 0.9 (0.79) | 10.7 (0.78) | 30.9 (0.97) |
| Pine, rich | CWTL | 18.2 (1.00) | 7.5 (0.97) | 11.9 (0.99) | 0.4 (0.54) | 11.3 (1.00) | 12.6 (1.00) |
| Open, poor | HIGH | 0.5 (0.56) | 0.0 (0.54) | 0.0 (0.42) | 0.0 (0.06) | 0.0 (0.50) | -0.8 (0.40) |
| Open, poor | CWTL | 2.1 (0.71) | 1.4 (0.64) | 0.5 (0.56) | -6.4 (0.05) | 6.9 (0.99) | 2.2 (0.81) |
| Open, rich | HIGH | 26.5 (0.93) | -19.4 (0.10) | 27.9 (1.00) | 0.0 (0.00) | 27.9 (1.00) | 53.6 (0.98) |
| Open, rich | CWTL | 12.5 (1.00) | 1.8 (0.77) | 19.3 (1.00) | 10.7 (1.00) | 8.5 (1.00) | 6.2 (1.00) |

Table S3. Changes and initial differences in taxa abundances in different peatland types. *Betula pubescens* could not be modelled in poor open mires due to low occurrence. Values are medians of 1 000 MCMC samples. Values inside parenthesis are proportion of samples that had positive sign for median.

| Peatland type | Taxon | Restoration effect, 10 yr | Initial difference, restored vs. drained | Change, restored | Change in restored vs. pristine, yr |
| --- | --- | --- | --- | --- | --- |
| Pine, Poor | Betula pubescens | 0.0 (0.74) | -0.0 (0.14) | -0.0 (0.02) | -0.0 (0.04) |
| Pine, Poor | Eriophorum vaginatum | -3.5 (0.00) | 2.0 (0.99) | -0.2 (0.42) | 3.5 (0.99) |
| Pine, Poor | Sphagnum spp. | 4.5 (0.82) | 16.3 (0.98) | 4.4 (0.85) | -2.3 (0.34) |
| Pine, Poor | Forest mosses | -6.4 (0.00) | 1.1 (0.69) | -1.9 (0.03) | -1.9 (0.02) |
| Pine, Poor | Carex spp. | 0.0 (0.55) | 0.0 (0.09) | 0.0 (0.48) | -0.0 (0.19) |
| Pine, Rich | Betula pubescens | 0.3 (0.99) | -0.0 (0.38) | 0.2 (0.97) | 0.2 (0.98) |
| Pine, Rich | Eriophorum vaginatum | 9.3 (1.00) | 0.9 (0.93) | 9.0 (1.00) | 9.0 (1.00) |
| Pine, Rich | Sphagnum spp. | 17.7 (1.00) | 8.6 (0.92) | 13.6 (1.00) | 0.5 (0.53) |
| Pine, Rich | Forest mosses | 0.4 (0.57) | -10.0 (0.01) | -1.6 (0.00) | -1.6 (0.00) |
| Pine, Rich | Carex spp. | 3.2 (1.00) | 0.3 (0.71) | 2.5 (1.00) | 1.7 (0.90) |
| Open, Poor | Betula pubescens | N/A | N/A | N/A | N/A |
| Open, Poor | Eriophorum vaginatum | -0.7 (0.38) | -1.0 (0.24) | 2.7 (0.94) | 2.0 (0.95) |
| Open, Poor | Sphagnum spp. | 11.0 (0.88) | -10.1 (0.22) | 17.7 (0.99) | 18.8 (0.94) |
| Open, Poor | Forest mosses | -0.3 (0.10) | -0.2 (0.06) | -0.0 (0.04) | -0.0 (0.04) |
| Open, Poor | Carex spp. | 0.0 (0.77) | 0.0 (0.17) | 0.0 (0.68) | 0.0 (0.56) |
| Open, Rich | Betula pubescens | 0.0 (0.70) | 0.0 (0.61) | 0.0 (0.40) | 0.0 (0.40) |
| Open, Rich | Eriophorum vaginatum | 4.4 (1.00) | 0.6 (0.76) | 4.8 (1.00) | 4.3 (1.00) |
| Open, Rich | Sphagnum spp. | 6.3 (0.88) | 1.9 (0.60) | 2.1 (0.7) | 7.6 (0.82) |
| Open, Rich | Forest mosses | -0.2 (0.32) | -0.7 (0.07) | -0.2 (0.00) | -0.2 (0.00) |
| Open, Rich | Carex spp. | 0.7 (1.00) | -0.0 (0.41) | 0.8 (1.00) | -0.5 (0.39) |

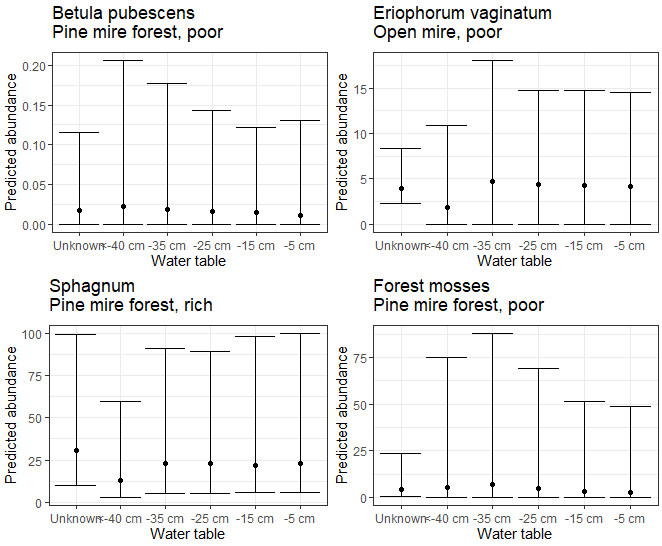

Figure S4. Predicted abundances of different taxa in different water table scenarios. Median of 1 000 MCMC samples are shown as dots and whiskers show highest and lowest 10 % of the samples.
